# Monalisa: an open source, documented MATLAB toolbox for magnetic resonance imaging reconstruction

**DOI:** 10.1101/2025.06.06.658299

**Authors:** Mauro Leidi, Yiwei Jia, Dominik Helbing, Jaime Barranco, Berk Can Açikgöz, Eva Peper, Jean-Baptiste Ledoux, Jessica A.M. Bastiaansen, Bastien Milani, Benedetta Franceschiello

## Abstract

**Purpose:** An open-source, user-friendly MATLAB framework for Magnetic Resonance Imaging (MRI) reconstruction was developed to simplify the reconstruction process, with a specific focus on non-Cartesian imaging and dynamic applications in the presence of motion.

**Methods:** Monalisa is decomposing the reconstruction pipeline into clear modular steps, including raw data reading with flexible file-type abstraction, trajectory computation, density compensation, advanced coil sensitivity mapping, and tailored binning strategies through its “mitosius” preprocessing stage. The framework supports a suite of reconstruction methods, including iterative-SENSE (also named CG-SENSE), GeneRalized Autocalibrating Partial Parallel Acquisition (GRAPPA) reconstructions, and regularized reconstructions supporting both spatial and temporal regularization using *l*_1_ (Compressed Sensing (CS)) and *l*_2_ techniques, accommodating both Cartesian and non-Cartesian acquisitions. We performed benchmark experiments comparing Monalisa with the Berkeley Advanced Reconstruction Toolbox (BART) toolbox on simulated 2D radial acquisitions.

**Results:** Results of the comparison demonstrate competitive performance, yielding higher Structural Similarity Index (SSIM) and lower *l*_2_ error. Notably, Monalisa reconstructions exhibited fewer visible artifacts than BART.

**Conclusion:** By providing comprehensive documentation, Monalisa serves not only as a powerful tool for research and clinical imaging but also as an educational platform to facilitate innovation in MRI reconstruction.

## 1 INTRODUCTION

MRI is a powerful, non-invasive, and versatile technique, considered a pillar in modern medical practice. It provides measurements that reflect properties of tissues, organs, and their pathologies ^1^, or changes in oxygen level in neural tissue, linked to neural activity, known as functional MRI ^2,3,4^. MRI enables the investigation of a wide variety of physiological phenomena, based on the physics of nuclear magnetic resonance ^1^: the properties of the nuclei are measured in the spatial-frequency domain, called k-space. Images can be created with contrasts reflecting tissue properties (determined by proton density, T1 and T2 relaxation times of the nuclei) or tissue susceptibility variations inducing magnetic field inhomogeneities. MRI measurements can also reflect biomechanical properties, electrical currents, oxygen levels, among other applications ^1^. The reconstruction process transforms measured k-space data into images that can be used for diagnostic or research purposes.

Acquisition and reconstruction strategies have evolved significantly to address the challenge of long acquisition times and motion sensitivity. A key approach to acceleration is partial sampling, which reduces the number of acquired data points but introduces aliasing artifacts that require correction during reconstruction ^5^. While full Cartesian sampling offers uniform k-space coverage, it is often impractical at high spatial resolution due to long scan times and its sensitivity to motion ^6^. Partial Cartesian sampling has been proposed ^5^, but this approach generates coherent aliasing artifacts that are difficult to correct. Alternatively, non-Cartesian schemes, such as radial acquisition schemes, introduce incoherent aliasing, which can be effectively mitigated using advanced reconstruction algorithms ^7,5^. Additionally, non-Cartesian sampling schemes—such as 2D golden-angle radial ^8,9^, 3D spiral phyllotaxis ^10^, and to some extent stack-of-stars ^11^ improve robustness to motion by repeatedly sampling the k-space central regions ^12,13,14^. Both partial and non-Cartesian sampling come at the expense of more complex and slower reconstruction techniques. While these schemes are used and enable high-quality image reconstructions, the theoretical framework underlying these methods is less fully established than for Cartesian sampling. In particular, the lack of a general sampling theorem does not allow to rigorously define key concepts such as a fully sampled k-space and effective spatial resolution. Moreover, the reconstruction of non-Cartesian or motion-resolved datasets is often not straightforward and can require substantial implementation effort. These challenges possibly limit broader adoption among researchers. K-space undersampling remains a challenge in MRI, especially when imaging moving organs such as the heart, lungs, and eyes. To address these challenges, several innovative computational techniques have been developed, each leveraging distinct aspects of the acquisition process:

- **Accelerated Parallel Imaging (PI):** SMASH ^15^ uses the spatial information provided by specific arrangement of surface coils to replace part of the phase encoding typically generated by magnetic field gradients, enabling partially parallel image acquisition. Techniques such as Sensitivity encoding (SENSE) ^16^, GRAPPA ^17^, and iterative-SENSE ^18^ exploit coil sensitivity profiles either explicitly (SENSE, iterative-SENSE), or implicitly (GRAPPA), to both shorten acquisition times and reduce undersampling artifacts. By sharing the spatial information provided by multiple receiver coils in order to fill in the missing data, these methods can reconstruct images from incomplete k-space data.
- **CS:** This approach is based on the premise that the image to be reconstructed is sparse in a certain domain (i.e., a vector basis) ^19^, and that the measurements are acquired in a randomized manner, leading to incoherent artifacts that resemble random noise. CS allows to share information between incompletely measured datasets and some prior knowledge. This allows to complete the missing data and enable high-quality reconstructions from reduced partially sampled datasets, thereby reducing scan times ^20,21,5,22,23^. For example, a strong prior knowledge in CS reconstructions is that temporal neighboring frames in a cinematic image (CINE) must be very similar. In that case, CS can efficiently be used to share information between different frames in order to complete the unacquired information. However, integration of CS into clinical workflows is still limited by its high computational demands and prolonged reconstruction times ^24^.
- **Deep Learning (DL):** Recent advances in DL have transformed MRI reconstruction by learning mappings between undersampled k-space and fully reconstructed images. The strategy of DL reconstruction is to share information between the incompletely acquired data and those acquired from other patients present in the training dataset, in order to complete the missing data. These “stronger together” methods have achieved significant improvements in both reconstruction speed and image quality, demonstrating notable potential for clinical applications ^25,26^. Most physics-based DL reconstruction methods developed so far rely on supervised learning, which requires high-quality datasets consisting of paired undersampled k-space data and corresponding ground truth images. These ground truth images are typically obtained by fully sampling and reconstructing the data. However, for high-dimensional imaging (4D and beyond), this is impractical, as acquiring even a single fully sampled MRI sequence can take several hours. As a result, training datasets for such high-dimensional data must be reconstructed using conventional non-DL methods. Recently, fully self-supervised approaches have emerged as a promising alternative ^27^, enabling networks to be trained with loss functions that do not rely on any reconstructed images. These methods have shown encouraging performance, although for the moment they remain limited to two-dimensional imaging.

While the theoretical framework for reconstruction is rooted in well-established mathematical principles, the practical implementation of these processes often remains opaque. On the one hand, this is justified, as MRI remains a clinical tool used in everyday diagnostic processes. On the other hand, the transition from raw data to images at the MRI console is often perceived by imaging researchers as a “black box”, with proprietary closed-source algorithms limiting accessibility. Recently, remarkable efforts have been made to open the “black box”: several open source initiatives have emerged ^28,29,30,31,32,33,34,35,36,37^, democratizing access to MRI reconstruction capabilities and fostering innovation across research and clinical settings. Among these, we highlight **BART**, a versatile toolkit supporting PI, non-Cartesian reconstruction, and machine learning integrations. BART provides an extensive suite of tools for advanced MRI reconstruction workflows and is widely adopted in the research community ^28^. **MRIReco.jl** ^29^ is another open source toolkit written in Julia, reusing existing functionality from other efficient Julia packages and optimized for MRI-related operations. MRIReco.jl’s modularity allows for efficient MRI reconstruction workflows. Reflecting this broader trend, vendors have begun to engage in similar efforts; for instance, Siemens Healthineers recently launched **OpenRecon**, a vendor-supported interface allowing integration of external software directly at the scanner console. Open-source reconstruction toolboxes still face limitations in usability and learnability. The inherent complexity of image reconstruction and the need for computationally efficient implementations force the adoption of low-level programming languages and modular frameworks, making the code difficult to digest. Additionally, the rapid introduction of new features to BART often outpaces documentation updates, potentially making it difficult for novice developers seeking to learn or contribute. In this article, we aim to introduce **Monalisa**, a comprehensive, open source toolbox for MRI reconstruction. Monalisa supports both Cartesian and non-Cartesian sampling, it supports advanced techniques such as CS and iterative reconstructions. **Monalisa is specifically designed to address the challenges of non-Cartesian and motion-resolved reconstruction** workflows, with the goal of simplifying their implementation and making them more accessible. Monalisa is MATLAB-based, easy to install, well documented, making it a good educational tool for learning and teaching MRI reconstruction.

## 2 METHODS

### 2.1 Notations and Definitions

To facilitate understanding, we introduce the following notation and definitions used throughout this work, to clarify the variables used in the implementation. For simplicity, the theoretical foundations of the reconstruction process are presented in the supplementary material section.

- **r**: is a vector whose components represent the spatial coordinates in the spatial domain. The spatial domain is the physical space, where the imaged object is located. These coordinates are defined with respect to the center of one voxel next to the center of the image, which serves as the origin for the reconstruction.
- **k**: is a vector whose components represent the coordinates of a spatial frequency in the k-space. These coordinates are defined with respect to the center k-space, which expresses spatial frequency of 0 (no oscillation in space). The reference axes in the spatial frequency domain are named *k*_*x*_, *k* _*y*_ and *k* _*z*_.
- **Frame**: a frame represents a single snapshot of the physical space—whether in 2D or 3D—much like a photograph. However, instead of containing conventional pixel intensities, each frame is the list of complex values assigned to each voxel-center of a Cartesian grid. In Monalisa, these frames are stored as arrays.
- *v or* **frDim**: the spatial dimension of a frame (*fr Dim* = 2 for 2D frames, *fr Dim* = 3 for 3D frames).
- **nFr**: the number of frames of a given image. For example, if the image is a sequence of 10 different temporal states, *nFr* would be 10. This is important since the definition of an image can differ from common usage: a single image can contain multiple frames, making it more akin to a video in everyday language.
- **Image**: an image is a ordered group of *nFr* complex-valued frames each of dimension *fr Dim*. In Monalisa, we store images using a cell array of size *nFr*, each cell containing a distinct frame.
- **frSize**: the frame size (commonly referred to as matrix size in the MRI community) defines the number of discrete samples along each dimension (*fr Dim*) within a given frame. For example, if we have *nFr* = 5 frames, each with *fr Dim* = 3 and *fr Size* = [120, 120, 120], it means that we are reconstructing five 3D complex frames, each with 120 × 120 × 120 voxels.
- **x**: any possible complex-valued anatomical image in the spatial domain.
- **x**^*true*^: the physical underlying complex-valued image. The ideal output if the measurements and reconstruction processes were perfect, assuming that the physical model is exact.
- **x**^*targ*^: the target complex-valued image. Since obtaining *x*^*true*^ is not feasible for reasons explained in the Coil Sensitivity section of the supplementary material, the reconstruction instead aims to recover an image *x*^*targ*^ that closely resembles *x*^*true*^ but is not identical to it.
- **x**^#^: the complex-valued reconstructed image in the spatial domain. This is simply the output of the reconstruction algorithm. A successful reconstruction yields *x*^#^ as close as possible to *x*^*targ*^.
- **x**_*l*_: the image obtained via the receiver coil number *l*, also named coil image number *l*, in the spatial domain for a given candidate image *x*.
- **y**: an element of the data-space. Hence it represent any possible outcome of an acquisition. We define *nSamp* as the total number of measured complex values. The elements of *y* are referred to as measurements or samples. The total number of k-space points along the acquisition trajectory is denoted as *nPt*. The number of coils or channels is represented by *nCh*. Hence, *nSamp* is equal to *nCh* times *nPt*. The sampled points are composed by readout lines, each containing *N* single trajectory points. In Monalisa, we store raw data using arrays of size [*nC h, N, nLines*] or a permutation of it.
- **Y**: the data space, a complex vector space that contains all possible vectors y.
- **y**_0_: the raw data acquired by the coils during the MRI experiment. As part of the reconstruction preparation, *y*_0_ can be modified by redistributing its samples among different bins and filtering out certain readouts. Even after preparation, the data remains labeled as *y*_0_ and is referred to as the reconstruction data.
- **C**_*l*_: : The (relative) coil sensitivity map for the *l*-th receiver coil. Its size at the time of estimation is *C*_ *fr Size*, while its dimension is *fr Dim*. At reconstruction time, *C*_*l*_ is interpolated so that its size becomes equal to *fr Size*.
- **C**: The coil sensitivity map. In MRI theory *C* is often defined as the matrix resulting from vertical concatenation of each single *C*_*l*_. In Monalisa implementation, *C* is estimated as an array of size [*C*_ *fr Size, nC h]*. Just before reconstruction, *C* is interpolated to a single precision array of size [ *fr Size, nC h*].
- 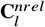 : The non-relative coil sensitivity map of the *l*-th receiver coil. This is usually referred to as coil sensitivity in the literature. For a given image *x*, the associated coil images *x*_*l*_ is given by 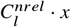.
- **t**: The k-space complete acquisition trajectory. The size of the trajectory is [ *fr Dim, nPt*]. As part of the reconstruction preparation, *t* can be modified by redistributing its samples among different bins and filtering out certain readouts. Even after preparation, the trajectory is still referred to as *t* and is referred to as the reconstruction trajectory.
- **B**_1_,… , **B**_*nB*_: The user-defined binning masks. *nB* is the number of binning masks, which is 1 or 2 for the reconstructions currently implemented in Monalisa. The size of binning mask number *b* is [*nBin*_*b*_, *nLines*], where *nBin*_*b*_ is the number of bins in binning mask number *b*.
- **nBins**: The total number of bins, or groups of data from which single frames are reconstructed. Since binning masks are combined using a Cartesian product, *nBins* = *nBin*_1_ · … · *nBin*_*nB*_ = *nFr* .
- **Chain**: A one dimensional array of cells (i.e. a Matlab one-dimensional cell-array). Every cell can contain a multi-dimensional arrays, such as a frame, a trajectory or a raw-data array.
- **Sheet**: A two dimensional array of cells.

### 2.2 Reconstruction Methods

This section provides a brief overview of the reconstruction methods available in Monalisa by presenting the corresponding reconstruction problems for each (except for GRAPPA reconstructions). Monalisa primarily supports non-Cartesian reconstructions, which rely on data gridding to handle non-uniform sampling of k-space. However, for many of these methods, an equivalent reconstruction is also implemented for Cartesian acquisitions, solving the same optimization problem without requiring a gridding step. These Cartesian versions can be identified by function names ending with the suffix “_partialCartesian”. An exception to this naming pattern is the non-Cartesian method `bmMathilda`, whose Cartesian counterpart is called `bmNasha_partialCartesian`. Despite the suffix, methods ending in “_partialCartesian” are not restricted to undersampled data; they can also be applied to fully sampled Cartesian acquisitions.

In general, a reconstruction algorithm aims to find an image estimate *x*^#^ that best explains the measured k-space data *y*_0_ given a forward model *A*, while also incorporating any available prior knowledge, if provided. This model *A* is the composition of the (Cartesian or non-Cartesian) spatial Fourier transform *F* and the coil sensitivities matrix *C* (see Coil Sensitivity section in supplementary material):

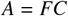

This model maps any candidate image *x* to the modeled data *Ax*. For non-regularized reconstructions, the goal is simply to minimize the distance between the measured data *y*_0_ and the modeled data *Ax*, where the distance is usually defined in terms of *l*_2_-norm ^38^:

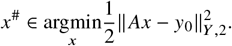

Regularization is a way to incorporate prior knowledge. It is done by adding a regularization term *R*(*x*) to the objective function ^38^. This term is balanced with the data fidelity term by a weighting factor *λ*:

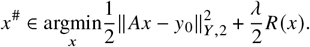

A well-designed regularization function *R* formalizes our prior knowledge by assigning low values to any image exhibiting the desired property. By including *R*(*x*) in the objective function, the reconstruction then favors only those candidates *x* that conform to what we already know about the true image, among all candidates for which the modeled data *Ax* is close to *y*_0_. For example, *R* may be a measure of the simplicity of *x*, in which case the associated prior knowledge is that the solution *x*^#^ we are looking for is sparse in a given vector basis. The theory of compressed sensing ^19,39,40^ specifies some conditions under which *R* can be written as a *l*_1_-norm. Monalisa’s reconstruction methods are the following:

#### A Zero-Padded, Gridded, Non-Iterative Reconstruction

(`bmMathilda`) for non-Cartesian sampling. It performs an approximative-inverse Fourier transform for each coil and, if the coil sensitivity matrix *C* is provided, combines the resulting coil images with the pseudo-inverse of the coil sensitivity matrix. This approximative-inverse is a linear map that we will write *F*^∼1^ so that

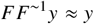

for any raw data vector *y*. The implementation of *F*^∼1^ is the composition of:

1. A gridding step, which is a discrete convolution finding an approximation of *y*_0_ on the positions of a Cartesian grid ^41,42^. The value of any position on the Cartesian grid that is not sufficiently close to *t* is set to 0 (zero-padding),
2. A conventional inverse discrete Fourier transform (DFT),
3. A deapodisation to cancel the filter resulting from the gridding operation.

Writing *F*^∼1^*y* the list of coil images, multiplying it by the coil sensitivity pseudo-inverse leads the solution of

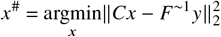

This is the result returned by Mathilda. It is by construction an approximate solution of the least-squares problem solved by Sensa (see hereafter) but it does not share the information between coils in order to fill the missing unsampled information, as it is done in SENSE and iterative-SENSE reconstructions. The Cartesian counterpart of Mathilda is Nasha_partialCartesian.

#### A 2D and a 3D GRAPPA implementation (bcaNeith2 and bcaNeith3)

for uniformly partially sampled Cartesian trajectories. Like Mathilda, it is a non-iterative single-frame reconstruction. However, it does not work directly with coil sensitivity maps. By its design, GRAPPA needs a calibration scan that consists of a few fully sampled lines acquired in the neighborhood of the k-space center. These few lines implicitly carry out information about the coil sensitivity maps and they are used to build some GRAPPA kernels, which are then convoluted with the measured data to complete the non-sampled part of k-space. Since this technique does not require an explicit estimation of the coil sensitivities, we say that it is using coil sensitivities implicitly. We refer the reader to the original manuscript for additional information ^17^.

#### An Iterative SENSE implementation (bmSensa) solving

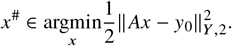

Sensa is our implementation of iterative-SENSE ^18^ using sensitivity encoding (SENSE) and iterative Conjugate Gradient Descent (CGD). This method is useful for both Cartesian and non-Cartesian data and can reconstruct images from incompletely sampled data by sharing information between different channels. However, it can struggle with heavily undersampled data because it operates frame-by-frame without sharing information between frames, making it less suitable for dynamic imaging. Despite this limitation, iterative-SENSE plays a fundamental role in the theoretical framework of image reconstruction.

#### A least-square reconstruction with *l*_2_ spatial regularization (bmSleva) solving

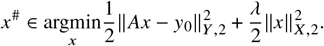

Sleva is a single-frame reconstruction method. It uses *l*_2_-regularization in order to make the solution unique. As *h* approaches zero, the optimization problem associated with Sleva returns the solution to the least-squares problem with smallest *l*_2_-norm described previously. Unlike the *l*_1_-norm, the *l*_2_-norm regularization does not enforce sparsity in any form. Sleva is therefore not a CS reconstruction.

#### A least-square reconstruction with *l*_1_ spatial regularization (bmSteva) solving

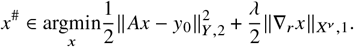

Steva is a single-frame reconstruction method. It is a CS method with spatial regularization by discrete anisotropic total variation. The *l*_1_-norm enforces sparsity in the spatial gradient of the image written ∇_*r*_ *x*, which means that it encourages many coefficients of ∇_*r*_ *x* to be exactly zero. This helps preserve sharp edges and fine details in spatially piecewise-constant images.

#### A least-square reconstruction with *l*_2_ temporal regularization, with or without motion compensation (bmSensitivaMorphosia_chain) solving

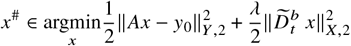

If deformation matrices are provided to the reconstruction function, then 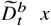 denotes the list of (backward) motion-compensated temporal differences, and it follows that the regularization is the sum of the squared *l*_2_-norms of the motion-compensated finite differences between temporal neighbors. Otherwise, 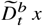 corresponds to the (backward) finite-difference time derivative of *x*, and the the regularization is the sum of the squared *l*_2_-norms of the finite differences between temporal neighbors. This reconstruction strategy is specifically designed for dynamic imaging applications.

#### A least-square reconstruction with *l*_1_ temporal regularization, with or without motion compensation (bmTevaMorphosia_chain) solving

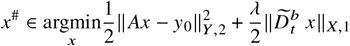

Again, if deformation matrices are provided to the reconstruction function, then 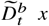 denotes the list of motion-compensated temporal differences. It follows that the *l*_1_-regularization enforces sparsity in the motion-compensated (backward) differences between temporal neighboring frames. Otherwise, 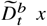 corresponds to the finite-difference time derivative of *x*, and the the *l*_1_-regularization enforces sparsity in the (backward) differences between temporal adjacent frames.

#### A least-square reconstruction with *l*_2_ backward and forward temporal regularization, with or without motion compensation (bmSensitivaDuoMorphosia_chain) solving

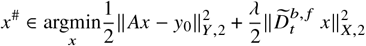

The SensitivaDuoMorphosia_chain method extends SensitivaMorphosia_chain by using both forward and backward finite-difference temporal derivatives.

#### A least-square reconstruction with *l*_1_ backward and forward temporal regularization, with or without motion compensation (bmTevaDuoMorphosia_chain) solving

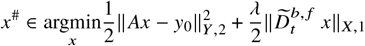

The TevaDuoMorphosia_chain method extends Teva-Morphosia_chain by using both forward and backward finite-difference temporal derivatives.

#### A least-square reconstruction with *l*_2_ temporal regularization, with or without motion compensation, with 2 temporal dimensions (bmSensitivaMorphosia_sheet) solving

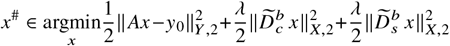

Here, the subscripts *c* and *s* correspond to the parameters of the two temporal dimensions. Note that by “temporal” we mean here any non-spatial dimension that can be interpreted as a time or a pseudo-time. Typically, *c* represents the cardiac time, while *s* denotes the respiratory time, meaning the time with respect to the start of the last cardiac or respiratory cycle respectively. The linear map 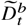 for *t* ∈ {*c, s*} have the same meaning as above for each pseudo-time: if deformation matrices are provided to the reconstruction function, then 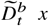 denotes the list of motion-compensated temporal differences between temporal neighbors. Otherwise, 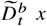 corresponds to the finite-difference pseudo-time derivative of *x*

#### A least-square reconstruction with *l*_1_ temporal regularization, with or without motion compensation, with 2 temporal dimensions (bmTevaMorphosia_sheet) solving

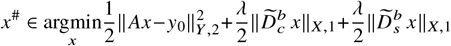

It is the *l*_1_ version of SensitivaMorphosia_sheet. Again, the linear maps 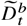 for *t* ∈ {*c, s*} have the same meaning as above for each (pseudo) time. .

For all reconstruction methods, the input arguments must include, among others, the content of the mitosius (y, t, ve, see Mitosius section in supplementary material) and the coil sensitivity matrix *C*. Additional reconstruction parameters are described in Table 1 .

**TABLE 1.** Parameter descriptions for reconstructions in Monalisa.

| Parameter | Description |
| --- | --- |
| **Coil sensitivity C** | Coil sensitivity maps as described in the Coil Sensitivity section of the supplementary material |
| **N<sub>u</sub>** | The size of the Cartesian grid used for regridding in k-space. It is independent from the frame size, but it is often chosen to match it. Reconstruction time depends on this parameter. Double-precision array with dimensions:<br>• $[N_x, N_y]$ for 2D spatial acquisitions<br>• $[N_x, N_y, N_z]$ for 3D spatial acquisitions |
| **dK<sub>u</sub>** | Step size of the grid used for regridding in k-space. Single-precision array with dimensions:<br>• $[dK_x, dK_y]$ for 2D spatial acquisitions<br>• $[dK_x, dK_y, dK_z]$ for 3D spatial acquisitions |
| **frSize** | Size of the reconstructed images in pixels. Double-precision array with dimensions:<br>• $[frN_x, frN_y]$ for 2D spatial acquisitions<br>• $[frN_x, frN_y, frN_z]$ for 3D spatial acquisitions |
| **Reconstruction Field of View (FOV)** | The reconstruction **FOV**, that can be different from the acquisition **FOV** for non Cartesian acquisitions (the acquisition **FOV** can be defined starting from the distance between two consecutive points along one line within the acquisition sampling pattern). |
| **ve_max** | Maximum volume elements value, an upper bound used to limit ve and avoid convergence issues. Single-precision scalar. |
| **nIter** | Number of iterations for the outer-loop of any iterative reconstruction. Double-precision scalar. |
| **nCGD** | Number of iterations for the **CGD** inner-loop to solve the least-square sub-problem of the ADMM algorithm. Double-precision scalar. |
| **x0** | Initial image for iterative reconstruction. Typically computed using a gridded reconstruction. Complex-valued, single-precision array. |
| **delta ($\delta$) or lambda ($\lambda$)** | Regularization parameter. Single-precision value, which can be:<br>• A scalar<br>• A list of two scalars $[\delta_{\min}, \delta_{\max}]$ where $\delta_{\min} < \delta_{\max}$<br>• A vector of length nIter |
| **rho ($\rho$)** | Convergence parameter of the **Alternating Direction Method of Multipliers (ADMM)** algorithm. Single-precision scalar. |
| **Gu** | The gridding (sparse) matrix used for forward gridding in iterative non-Cartesian reconstructions. |
| **Gut** | The transposed matrix of Gu used for backward (not inverse) gridding in iterative non-Cartesian reconstructions. |
| **Tu** | The deformation (sparse) matrix used for forward deformation in iterative temporally regularized reconstructions. |
| **Tut** | The transposed matrix of Tu used for backward (not inverse) deformation in iterative temporally regularized reconstructions. |

The output of the reconstruction process consists of one image (single-frame or multi-frame) in the spatial domain.

In some cases, providing **deformation matrices** (written *T*_*u*_ and *T*_*ut*_ for their transpose) to the reconstruction function improves the effect of a temporal (or pseudo-temporal) regularization in the presence of spatial misalignment between frames. Deformation matrices are used to compensate for motion. This can be particularly useful in applications like cardiac, kidney, liver, or lung imaging, where large motion occurs between frames. Note that the non-spatial dimension can be different from actual physical time. It can represent any list of motion states.

Each reconstruction method requires adjustment of the number of iterations and regularization parameters among others. The choice of these parameters depends on the degree of undersampling, the noise level, and the desired temporal resolution among other factors. In practice, a series of tests may be necessary to optimize these settings for a given dataset.

### 2.3 Comparison with existing reconstruction frameworks

To evaluate the performance of Monalisa compared to existing reconstruction frameworks, we conducted a controlled experiment comparing Monalisa and BART ^28^. The comparison was performed on three different 2D images, including a numerical Shepp–Logan brain phantom ^43^, a brain slice centered on the eye, and a cardiac slice (see Figure 1 a, ground truth column). The raw MRI measurements were synthetically generated by simulating a 2D radial acquisition with 30 lines of 512 points each, and the coil sensitivites associated with the images. For the Shepp-Logan phantom a simulated coil sensitivity was used. The trajectory was constructed by rotating each line sequentially by an angle of 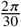 radians. The number of lines was deliberately chosen to induce undersampling artifacts in the gridded reconstruction, thereby highlighting the advantages of regularized reconstruction methods. Monalisa and BART were tested using iterative reconstructions with *l*_1_ and *l*_2_ regularization terms. For the *l*_1_-regularization case, we employed a total variation constraint to promote piecewise constant reconstructed images. For the *l*_2_-regularization case, we used the standard squared *l*_2_ norm of the image as constraint. The same coil sensitivity maps, trajectory and volume elements were used for both frameworks. Thus, the goal is to compare two implementations of the same reconstruction algorithm, using identical inputs, in order to demonstrate that our implementation quality is comparable to the state-of-the-art of open-source MRI reconstruction. Since reconstruction regularization can alter the reconstructed image support, a direct comparison of reconstructed and reference images can be misleading if evaluated on their raw intensity scales. To ensure fair evaluation, we applied an affine intensity alignment of the ground truth image magnitude to each reconstruction magnitude prior to computing similarity metrics. Specifically, for each reconstructed image x we estimated coefficients *a, b* by least-squares regression:

**FIGURE 1.**
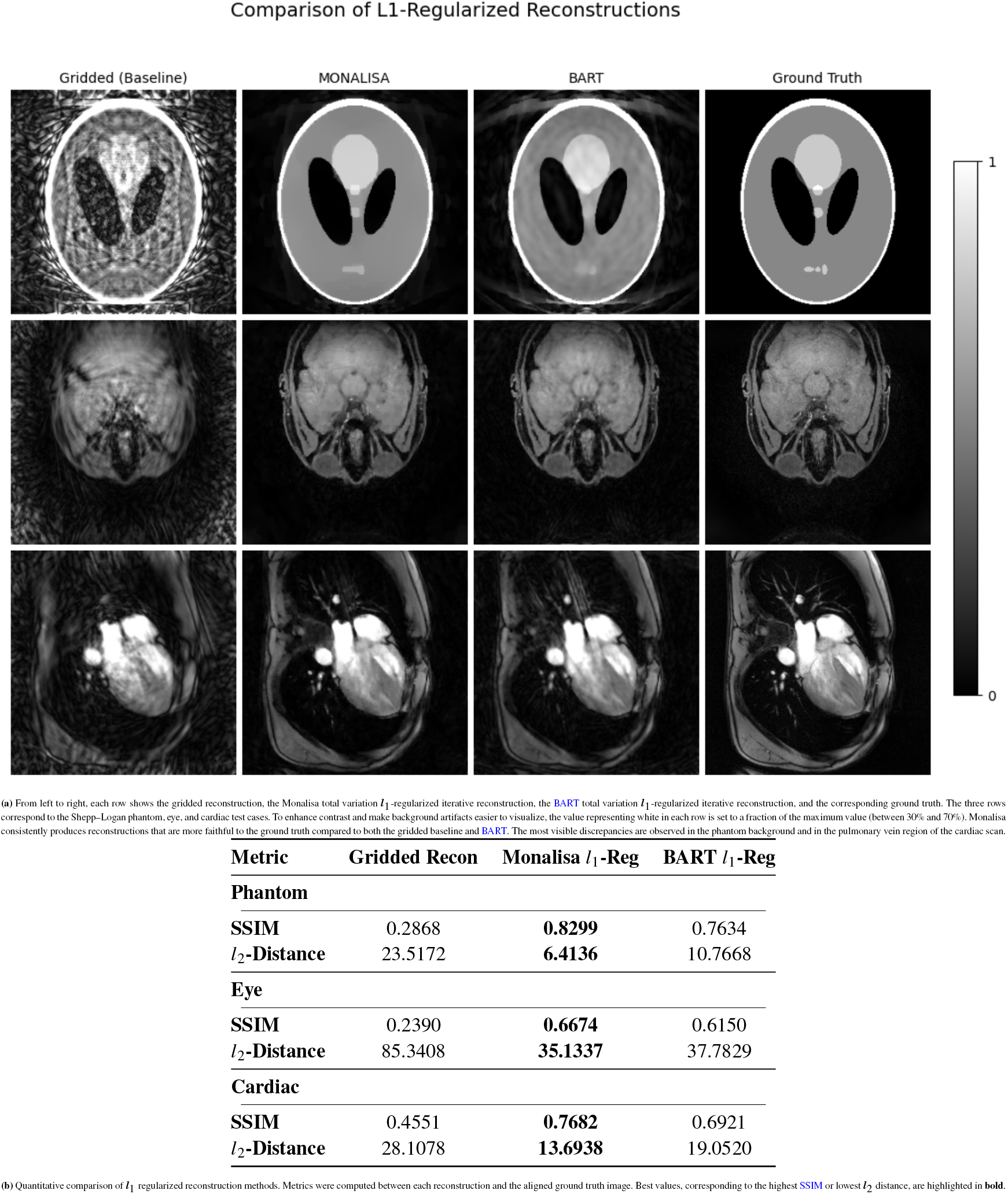
Comparison of *l*_1_-regularized reconstruction results using visual and quantitative metrics.

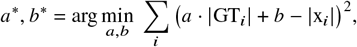

where *i* indexes all voxels in the images. The resulting aligned ground truth magnitude is

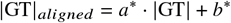

This alignment preserves the structural information in the images while compensating for arbitrary global scaling and offset differences between reconstructions and reference. SSIM and *l*_2_ distance were then calculated between the reconstructed magnitude images and the aligned ground truth magnitudes. For simplicity, in the remainder of this manuscript we refer to these aligned metrics simply as SSIM and *l*_2_ distance. The rationale for this step is that SSIM and *l*_2_ distance metrics are sensitive to shift in the reconstructed image support. A global linear transformation of magnitudes does not change relative contrast or structure, which are the quantities of diagnostic relevance and what human observers actually interpret. By bringing the ground truth onto the same photometric scale as the reconstruction, the evaluation reflects differences in structural fidelity rather than arbitrary scaling factors introduced by the reconstruction process.

To ensure fair evaluation, for both frameworks, regularization parameters were optimized via a grid search, and the regularization values yielding the highest SSIM were selected for each reconstruction independently. Reconstructions were performed using 160 iterations, a number which guarantees convergence for both frameworks. The number 160 was empirically determined by trying several incremental values and checking the convergence. For both frameworks, *l*_1_-regularized reconstruction was solved using ADMM, while *l*_2_-regularized reconstruction was solved using CGD. The maximum number of conjugate gradient iterations for ADMM was set to 4.

Reconstructions were evaluated using both SSIM ^44^ and *l*_2_ distance with respect to the associated aligned ground truth magnitudes. *l*_2_ distance computes the sum of squared differences between corresponding pixel intensities in two images. It is a straightforward, pixel-wise error measure of the reconstruction that treats every pixel equally. SSIM, on the other hand, is designed to mimic the human visual system by considering perceptual phenomena. It evaluates image quality based on three key components: luminance, contrast, and structure.

## 3 RESULTS

Images reconstructed using *l*_1_ and *l*_2_ regularization using the optimized regularization value are shown in Figure 1 a and Figure 2 a, respectively. To enhance contrast and make background artifacts easier to visualize, the value representing white in each row is set to a fraction of the maximum value (between 30% and 70%). Corresponding quantitative metrics are reported in Figure 1 b and Figure 2 b. Both the visual inspection and the metrics indicate that regularized iterative reconstructions outperform Monalisa’s gridded reconstruction across both frameworks.

**FIGURE 2.**
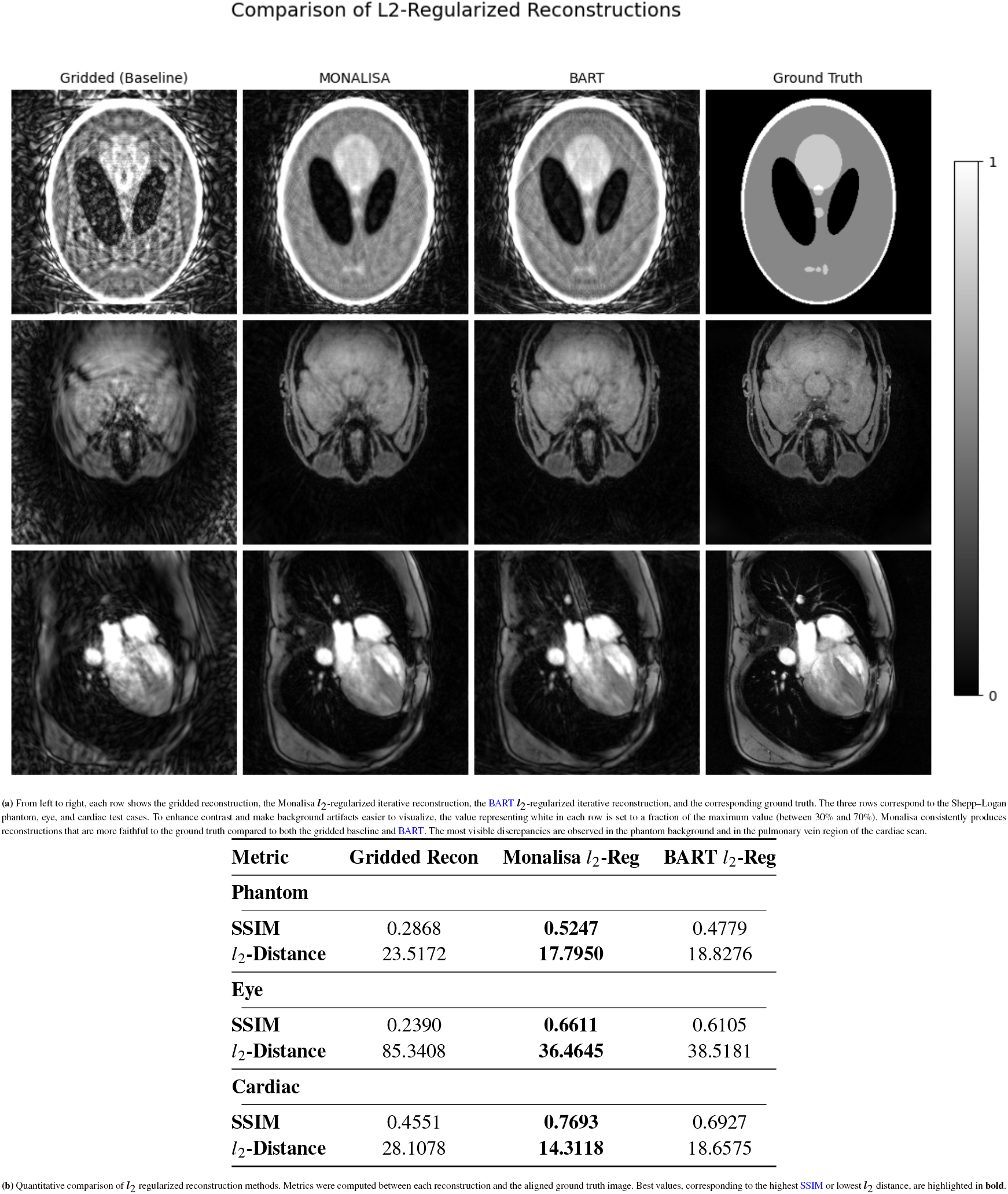
Comparison of *l*_2_-regularized reconstruction results using visual and quantitative metrics.

For both *l*_1_ and *l*_2_ iterative reconstructions, on all the three test examples, Monalisa achieves higher SSIM values and lower *l*_2_ distances. Visually, for the cardiac and phantom images, BART reconstructions appear more affected by artifacts, especially visible in the background for the phantom and in the pulmonary vein region of the cardiac image.

## 4 DISCUSSION AND CONCLUSIONS

In this study, we have presented Monalisa, a comprehensive and open source MRI reconstruction framework. We detail both software design choices behind Monalisa, and the steps required to transform raw k-space data into highquality images. By modularizing the reconstruction workflow, composed of the following steps: data reading, trajectory definition, volume elements computation, coil sensitivity estimation, binning, and mitosius preparation, Monalisa enables users to implement and experiment with a variety of reconstruction algorithms, including GRAPPA, iterative SENSE, CS with a variety of spatial and temporal regularization.

Our benchmarking against BART on simulated undersampled 2D radial data demonstrates that Monalisa achieves consistently better performance across the three evaluated test cases, both for *l*_2_-norm regularization and *l*_1_ total variation regularization. The observed differences might be attributable to variations in algorithmic implementation. However, the larger discrepancies observed in the phantom reconstruction background and in the cardiac pulmonary vein region remain unexpected, and their underlying causes have yet to be determined. A potential explanation is that BART’s generalized reconstruction options used are still experimental, as noted in BART’s pics helper documentation.

A key strength of Monalisa lies in its moderate complexity, thanks to its reliance on a high-level programming language and its documentation, which make the framework significantly simpler to understand. This accessibility makes Monalisa a great platform for teaching and learning MRI reconstruction, while also offering researchers an agile environment to test and develop new reconstruction strategies. Additionally, our framework directly integrates the recent trends of open source acquisition frameworks, supporting ISMRMRD raw data format ^45^, and trajectory generation with PulSeq ^46^. In doing so, Monalisa makes a significant contribution to the MRI community by promoting at the same time transparency and innovation.

However, Monalisa’s current implementation does not support specialized hardware acceleration (e.g., GPUs), which could enhance reconstruction speed, especially for large datasets. Nonetheless, this can be easily achieved by replacing the Fast Fourier Transform (FFT) operations with a CUDA-based alternative, which is called CUFFT (CUDA-based FFT implementation), or another GPU-based FFT implementation, as successfully demonstrated in previous work ^47,48^. Additionally, because Monalisa is built on MATLAB, a licensed environment, it may present accessibility challenges. Future work will focus on implementing GPU support and exploring more accessible development platforms to further improve both performance and accessibility.

Check out the documentation at: https://mattechlab.github.io/monalisa/

Check out the source code at: https://github.com/MattechLab/monalisa/

## Author contributions

**Conceptualization** and **Methodology** were carried out by M.L., B.M., and B.F.

**Software** development, including the design and implementation of reconstruction algorithms, coil-sensitivity estimation, gridding, volume elements computation, and related modules, was led by B.M., author of the first version of the software, with subsequent contributions from M.L. (Raw Data Reading, Dockerfile, compilation scripts), Y.J. (example scripts, code checkers, Dockerfile), D.H. (ISMRMRD support, inline documentation, UI for coil-sensitivity estimation), J.B., and B.C.A. (GRAPPA reconstructions). The design and implementation of the documentation were led by M.L., including infrastructure setup and most content contributions, with contributions from Y.J. (*Mitosius* section), B.M. (reconstruction section), and J.B. (auto-compiled documentation section).

**Formal Analysis** and **Visualization** were carried out by M.L., B.M., J.B., Y.J., and B.F.

**Investigation** activities, including data acquisition and preparation, were supported by J.B.L., E.P., and J.Bas., who also provided essential **Resources** and information on scanner hardware and acquisition protocols.

**Data Curation** and organization were supported by J.B., E.P., and J.B.L.

**Writing – Original Draft** was led by M.L., with significant contributions from B.M. (coil-sensitivity estimation and reconstruction sections) and B.F. (introduction).

**Writing – Review & Editing** and methodological **Validation** were performed by all authors.

**Scientific Supervision** was provided by B.F. and B.M.

**Project Administration** and **Funding Acquisition** were provided by B.F.

## Data Availability Statement

All benchmarking code and data presented in the Results are available at: https://github.com/MattechLab/comparisonMonalisaBart/. The software versions used are included as Git submodules in the repository and are therefore identifiable via the commit hashes visible in the repository.

## Acknowledgments

We thank all contributors and organizations whose support has made this project possible. All scientific content and interpretations are the authors’ own. This work was supported by the Swiss National Science Foundation (SNSF) (Grants: #220433, #229214 to Prof. Franceschiello; #320030B_201292 to Prof. Stuber; #320030B_194296 to Prof. Bastiaansen). The Sense Innovation and Research Center (Grant: KiCk fMRI, PI: Prof. Franceschiello & Prof. Dr. Juliane Schneider).

ChatGPT was used during the preparation of this manuscript for language editing and proofreading, and in limited cases for code-related assistance, primarily for minor refactoring and documentation generation. These tools were not used for scientific content generation, data analysis, or interpretation of results.

## SUPPLEMENTARY MATERIAL

### Monalisa workflow

Figure 1 illustrates the pipeline of Monalisa. Reconstructing raw data is a multi-step process combining trajectory definition, estimation of volume elements, coil sensitivity estimation, binning, mitosius preparation, and reconstruction. In this section, we describe all the steps that are necessary to transform raw k-space data into the desired reconstructed images.

#### 0.0.1 Reading Raw Data

Prior to any reconstruction or analysis, k-space raw data must be parsed into MATLAB objects. Monalisa achieves raw data-type abstraction using the RawDataReader ^∗^ interface class, an abstract base class that defines two key methods:

- **readMetaData**: this method is responsible for parsing header information and acquisition parameters from the raw data file. This includes extracting details such as the number of coils, acquisition lines, echo counts, and Field of View (FOV). Additionally, it computes other metrics, such as determining when steady state is reached by analyzing signal statistics.
- **readRawData**: this second method handles the extraction of the actual k-space data from the raw file. This method not only reads the data but also reshapes it into the expected MATLAB array format (typically organizing dimensions such as coils, readout points, number of acquisition lines). Moreover, depending on the provided flags, it can optionally filter out unwanted samples (e.g., non-steady-state lines)

Together, these two methods encapsulate the specifics of the data format, allowing the remainder of the Monalisa framework to interact with a uniform MATLAB object interface regardless of the underlying raw data file structure. Monalisa implements two subclasses that override these methods: one for Siemens raw data files^†^ and another for the ISMRM raw data format^‡ 1^. For the implementation of our Siemens raw-DataReader, we rely on mapVBVD^§^ originally developed by Philipp Ehses. Users working with a different file format must either convert their data to the ISMRM raw data format, or implement a new subclass that inherits from the base class.

#### 0.0.2 Trajectory

The raw data does not contain information about k-space trajectories, it solely contain *nSamp* k-space complex-valued measurements. Therefore it is crucial for the reconstruction process to know the precise spatial k-space location at which each measurement was acquired. In Monalisa, the trajectory is represented as a double-precision array **t** of size [frDim, nPt]; hence, each column encodes the k-space coordinates for one measurement, and each row encodes the k-space coordinates along one dimension. Our convention is that every acquisition trajectory must be defined in physical units derived from the scanner’s true input FOV^¶^ . In practice, this means that the k-space coordinates are expressed in physical units (e.g., mm^–1^), ensuring that the step size in each direction is directly linked to the intended FOV. For instance, if a Cartesian acquisition is designed for a FOV of [200, 300] mm, the step size along the *k*_*x*_ axis must be 1/200 (in mm^–1^) and along the *k*_*y*_ axis 1/300 (in mm^–1^). Similarly, for a radial trajectory with a FOV of [360, 360] mm, the spacing between consecutive points along each radial spoke must be 1/360 (in mm^–1^). If the trajectory has been computed using a different convention, the coordinates should be converted by applying the appropriate scaling factors, as outlined in the provided documentation^#^.

Monalisa provides a selector function, `t = bmTraj(mriAcquisitionNode)`, which offers several predefined k-space trajectories, including one that extracts trajectory coordinates directly from a PulSeq .seq file ^2^. It is important to note that when trajectories are generated without relying on a .seq file, the specific sampling points often depend on the implementation. Consequently, even nominally identical trajectories can produce slightly different coordinate sets across different sequence implementations. Therefore, it is crucial to ensure that the trajectory used during reconstruction corresponds exactly to the one intended during data acquisition.

#### Estimation of Volume Elements

Volume elements serve as a weighting mechanism that is essential for non-Cartesian Magnetic Resonance Imaging (MRI). The following qualitative explanation is intended to help the reader understand why these volume elements are useful. Imagine we want to compute the integral of a (well-behaved) integrable function *ƒ* over a 3D domain (e.g., 3D k-space):

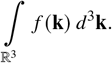

**FIGURE 1.**
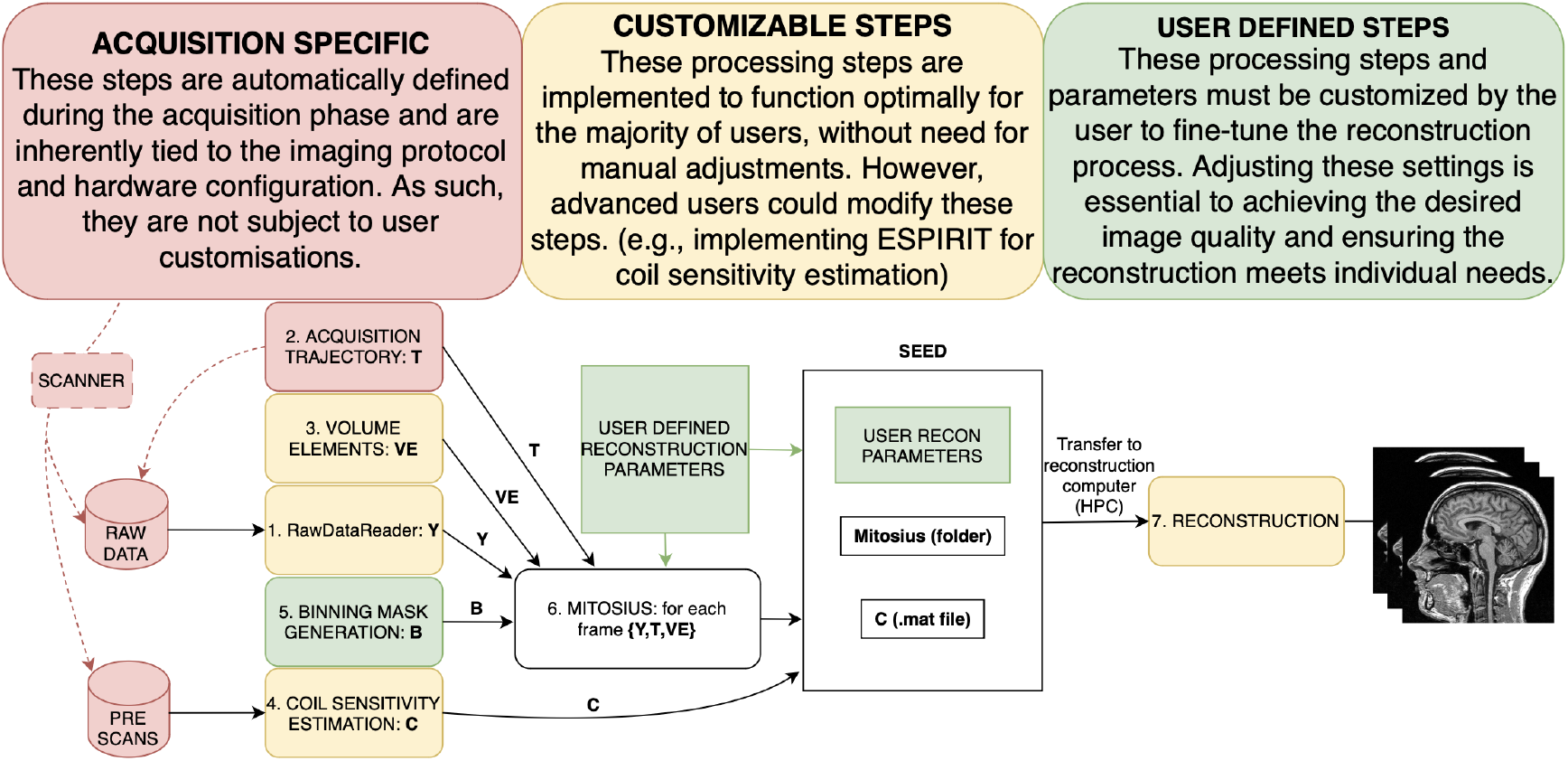
Simplified schematic of the pipeline in Monalisa. Information from the acquisition raw data files is processed through multiple stages, together with additional information, such as the trajectory, volume elements, and user-defined binning and reconstruction parameters. Colored blocks indicate the degree of user control: red denotes acquisition-specific steps that are fixed and cannot be modified after acquisition; yellow highlights configurable steps that are rarely changed but can be customized or improved; green represents user-defined steps that must be adapted for each reconstruction. Initially, the workflow defines binning masks alongside coil sensitivity estimation and volume element computation. This is followed by the mitosius preparation, and finally, image reconstruction. Refer to the reconstruction parameters table in the manuscript for a description of the reconstruction parameters.

In practice, on a computer, we must discretize this integral. If we know *ƒ* at a set of locations 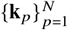, we can approximate the integral as:

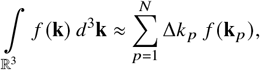

where Δ*k*_*p*_ is the volume element associated with **k**_*p*_. Now, consider that these sampling points {**k**_*p*_} belong to a trajectory **t**. If **t** is Cartesian, all points are equidistant and all the volume elements Δ*k*_*p*_ are identical. In this case, we have:

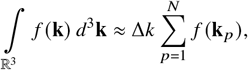

so that the influence of the volume elements reduces to a constant factor Δ*k*, which in fact can be ignored in MRI reconstructions after normalization. In contrast, for a non-Cartesian trajectory **t**, the volume element Δ*k*_*p*_ varies with the local point density. For example, radial trajectories oversample the k-space center, and for this reason the Δ*k*_*p*_ associated with those central points are lower. This non-uniformity requires each sample’s contribution to be weighted appropriately by its individual volume element. Failing to account for this variation can lead to artifacts in the reconstructed image and may compromise the convergence of iterative reconstruction algorithms. Intuitively, given a finite list of sampling points (i.e., the trajectory **t**), the volume element assigned to each point represents the volume “occupied” by that point relative to its neighbors. The local density is defined as the number of sampling points per unit volume, typically averaged over a small neighborhood (for example, within a spherical region). From this perspective, the local density at a given point is essentially the inverse of its associated volume element:

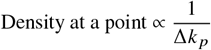

That is the reason why, historically, this concept is referred to as “density compensation” in MRI, originating from the transition from uniform trajectories, where the density is constant and can be neglected, to non-uniform trajectories. Although non-uniform density was once viewed as a problem to be “compensated”, it is in fact the general case, with uniform sampling being a special scenario ^∥^.

In Monalisa, volume elements are typically estimated as the volumes of the Voronoi cells associated with the trajectory points, where each Voronoi cell is defined as the region of space closer to a given point than to any other ^4^. It is important to note that these volume elements depend on the binning (see subsubsection 0.0.6)—since the density may change depending on which measurements are selected for a given frame—and are stored in a double-precision array **ve** of size [1, *nPt*]. No additional rescaling is applied, so the volume elements represent the absolute volume of the Voronoi cells.

In Monalisa, the function implemented to compute the volume elements of any reconstruction trajectory is `bmVolumeElement`. It is invoked using the following syntax:

`ve = bmVolumeElement(trajectory, type_of_trajectory, optional_arguments)`

This function acts as a selector based on the specified trajectory type. Users are encouraged to visit our documentation^∗∗^ to determine which trajectory type best suits their specific case. For any radial trajectory, we suggest using Voronoi based methods. Regardless of the adopted method, the user must define an upper bound `ve_max` for the volume elements, in order to avoid convergence issues in iterative reconstructions. As a rule of thumb, you can use `ve_max` to be a small multiple of the volume of a cell of the Cartesian k-space grid used for regridding (between 3 and 10). As an example,

for 2D frames :`ve_max = 10 · dKu_x · dKu_y`

for 3D frames :`ve_max = 10 · dKu_x · dKu_y · dKu_z`

#### 0.0.4 Coil Sensitivity Map Estimation

Parallel imaging employs arrays of many small receiver coils placed in close proximity to the tissues of interest. This close placement enhances the signal intensity detected by the coils, resulting in a high Signal-To-Noise Ratio (SNR) in the final image. However, this high SNR comes at a cost, which can be understood as follows. During acquisition, each region of tissue containing resonating nuclei acts as a source of electromagnetic waves that propagate through space, inducing a time-varying voltage in each coil according to the law of induction. The voltage measured by a given coil is the sum of contributions from all emitting sources, with each source’s contribution depending on its spatial location with respect to each part of the coil. This spatial dependence can be characterized by a complex-valued function, referred to as “coil sensitivity”, which describes how a given tissue region influences the voltage induced in a particular coil. Since coils for parallel imaging are small and very close to tissues, their spatial dependence is high, which means that the coil sensitivity of each individual proximity coil is spatially inhomogeneous. Moreover, the coil sensitivity of different coils is very different. Conceptually, we can say that each coil “sees” a very different image. The cost of using many surface coils resides therefore in the reconstruction challenge of combining these coil images into a single high-SNR image, a relatively complex process that requires accurate estimation of coil sensitivities. Coil sensitivity estimation can be either explicit or implicit, depending on the reconstruction method. One of the most widely used approach in the clinic is implicit estimation as part of GeneRalized Autocalibrating Partial Parallel Acquisition (GRAPPA) reconstructions, but all iterative reconstructions used in research need an explicit estimation, which often involves acquiring an additional calibration scan—a low-resolution, full-field-of-view image, used to compute coil sensitivities. This section presents *Sensitivity Iterative Gradient-descent Mapping Algorithm (SIGMA)*, a method for explicitly estimating coil sensitivities. Although this method was previously evaluated on patient data ^5^, the theoretical foundations and implementation details were not disclosed. In this work, we present a comprehensive development of both the underlying formalism and its practical implementation. *SIGMA* builds upon key ideas from an early proposal by Pruessmann ^6^ and shares some concepts with JSENSE ^7^.

We begin by detailing the coil sensitivity estimation method proposed in ^6^, as it closely resembles our approach. Given the ground truth image *G*^*true*^, the relationship between its coil images number *l* , 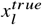, and the the coil sensitivity number *l*, 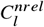, is a point-wise multiplication at each spatial location:

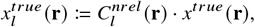

where 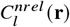 represents the coil sensitivity of coil *l* evaluated in position **r** and is a complex number. A higher magnitude of 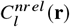 indicates a greater sensitivity of coil *l* at location **r**.

In ^6^, the first step in coil sensitivity estimation involves acquiring and reconstructing full-FOV images for each surface coil, denoted as 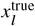, as well as an image from the body coil, denoted as 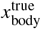. To perform an estimation of 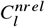, in ^6^ 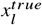 is then divided by 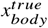. While not stated in the study, this method implicitly assumes that the sensitivity of the body coil is constant over space (homogeneous) in the Region of Interest (ROI). In fact, if that assumption is true, the body-coil image is given by

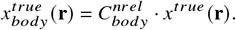

Note that here 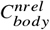 is independent of *r*. It holds then

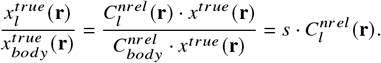

so that 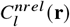 can be estimated by 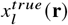 and 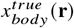 up to a scalar factor *s*. As a second step, to prevent the noise present in the images from altering the sensitivity estimation, ^6^ applies smoothing via polynomial fitting on a binary mask applied to the previously computed 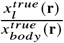 . To create this binary mask, a threshold is first manually computed to split noise from signal-containing region, where voxels with value smaller than the threshold are considered noise. Additionally, residual voxels that do have a sparse neighborhood are also considered noise. In a final step, an extrapolation zone is computed using a region growing around the mask. All voxels outside the mask and the extrapolation zone are assumed to be zero-valued, and polynomial fitting yields the final sensitivity estimates. We interpret this extrapolation zone as a heuristic designed to extend the sensitivity estimation slightly beyond the mask, to avoid abrupt changes in sensitivity at the edges. This helps reduce the oscillations often observed near discontinuities, particularly in polynomial fitting methods, which might lead to numerical inaccuracy or instability ^8^.

Our method SIGMA differs from the method of ^6^ primarily in four ways:

- We take advantage of the presence of two pairs of body-coils, instead of only using one. In conventional MRI scanners, there are usually *nBody* = 2 body coils.
- We achieve smoothing by a local voxel averaging, meaning that the value of a single voxels are replaced with the average of their value and those of all the adjacent voxels, instead of polynomial fitting.
- We extend the coil sensitivity estimate in regions without signal by solving a Laplace boundary value problem, instead of using the extrapolation zone method.
- We estimate the relative coil sensitivity maps, instead of non-relative coil sensitivity maps. By “relative” or “biased” coil sensitivity of channel number *l* we mean the coil sensitivity defined by

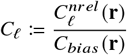

where *C*_*bias*_ (**r**) is any complex valued function of space that makes *C*_*ℓ*_ estimable in practice. This choice of estimating a relative coil sensitivity instead of the true non-relative coil sensitivity allows us to relax the assumption done in ^6^, which was to consider 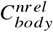 homogeneous in space. We will choose *C*_*bias*_ (**r**) proportional to the complex phase of one of the two body-coil sensitivities. Independently from our choice of *C*_*bias*_ (**r**), the reconstructed image using the maps *C*_*ℓ*_ as coil sensitivities is the image *x*^*targ*^ given by

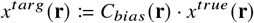

This follows from the signal equation ^9^

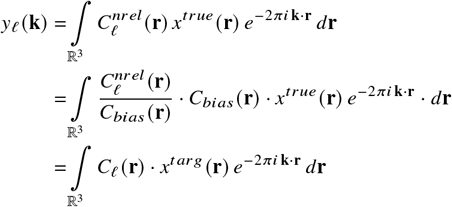

We are now ready to present the formalism and implementation of SIGMA. In order to estimate the maps *C*_*ℓ*_ in a region where resonating nuclei are located , we need the following definitions:

- the coil image number *l* of *x*^*targ*^

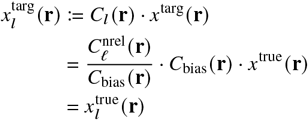

so that 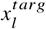 and 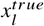 coincide, in contrast to *x*^*targ*^ and *x*^*true*^.
- The root-mean-squared body-coil image:

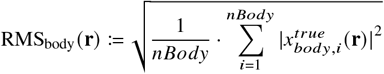
- The reference coil sensitivity:

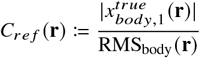
- The reference image:

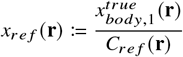

These three quantities can be estimated by only using coil images. We now present the following claim:

*Claim 1*. If it holds the approximation

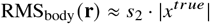

for some constant *s*_2_, and if we define

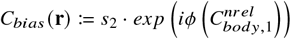

using *ϕ* (*z*) as the phase of the complex number *z*, and that scalar value *s*_2_, then it holds the approximation

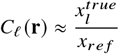

*Proof*. From the definitions and the assumption in **claim 1**, it follows that

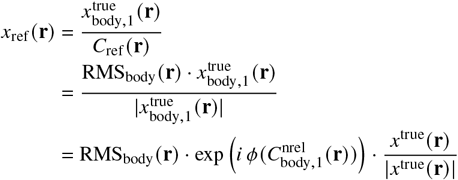

where *ϕ* (*z*) is the phase of the complex number *z*. Therefore

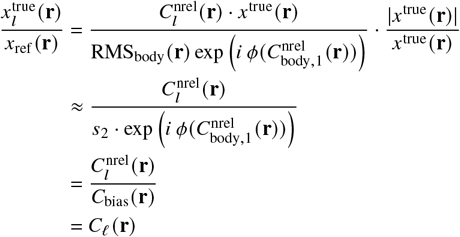

Since 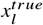 is easy to estimate, we can by **claim 1**, under the assumption that the Root Mean Square (RMS) of the body coil is proportional to the magnitude of *x*^*true*^, estimate *C*_*ℓ*_ in all locations where the resonating nuclei generate a signal significantly larger than the noise. Our assumption is weaker than assuming the existence of a single spatially homogeneous body coil, since we are only assuming we can estimate the magnitude of such a homogeneous coil, which in our case is a virtual coil that is obtained by combining both body coils.

We can now formulate our SIGMA method as the following steps:

- We estimate a binary mask that indicates which spatial voxels contain anatomical regions. Only the voxels that are within this estimated area will be used to estimate the coil sensitivity. This is justified, as the image outside the anatomical region should in theory be zero. The binary mask is generated through a thresholding process applied to both the Maximum Intensity Projection (MIP) ^10^ and RMS ^11^ images. Both MIP and RMS are methods to combine multiple surface coil images into a single representation. MIP takes the maximum intensity value across multiple-coil images, emphasizing bright structures and enhancing contrast-noise ratio ^12^. RMS uses a sum of squares combination, averaging out noise fluctuations ^13^. We implemented both manual and automated mask extraction. In manual mode, the user first visually adjusts a threshold on the MIP image, denoted as *mipthrs*, and on the RMS image denoted as *rmsthrs*. Additionally, a bounding box around the ROI is manually adjusted. In automatic mode, a threshold detection algorithm is implemented by fitting a bimodal Gaussian distribution to the histogram of pixel intensities for both images (RMS and MIP). The two Gaussian components are interpreted as representing the noise and signal distributions, respectively. The threshold is then set at the intensity value where the probability of the pixel belonging to the noise distribution equals the probability of it belonging to the signal distribution. Any pixel with value lower than the threshold is excluded from the mask. The automatic bounding box extraction works by first reshaping the 3D image and computing summed projections along each dimension. These projections are then normalized and binarized using Otsu’s method ^14^, allowing to isolate the largest connected component in each 2D projection. The bounding boxes for these components are determined and then merged—with added padding—to generate the final bounding box around the region of interest. For manual and automatic modes, the mask is then created by including only those voxels where the MIP value exceeds *mipthrs*, the RMS value exceeds *rmsthrs*, and that lie within the bounding box.
- We estimate the maps RMS_body_, *C*_*reƒ*_ and *x*_*reƒ*_ following the previous definition. We then estimate each relative coil sensitivity *C*_*ℓ*_ by **Claim 1** using the associated assumption. This leads to a first estimate of the coil sensitivities inside the mask.
- In order to mitigate the effect of noise on the estimation of each *C*_*ℓ*_ , we average the value of each voxel with the values of its neighbors, without considering voxels outside the mask to avoid issues at the boundary of the mask. This process achieves smoothing of the estimated values within the mask. For clarity we denote the smooth coil sensitivities as:

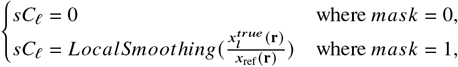
- Finally, to avoid abrupt changes in estimated coil sensitivity at the edges of the ROI, we extend the estimation outside the mask solving, via the Jacobi method ^15^, the Laplace boundary value problem. We denote the extended smooth coil sensitivity as *esC*_*ℓ*_ , obtained by solving two independent Laplace equations for the real and imaginary components:

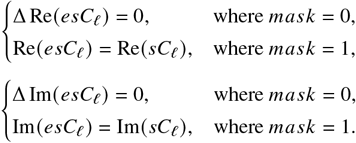

Here, Δ is the discrete Laplacian operator, and the smooth sensitivity values *sC*_*ℓ*_ inside the mask serve as Dirichlet boundary conditions.

Monalisa offers two implementation approaches for *SIGMA*: (1) using body coil calibration images and (2) using calibration raw data from which the corresponding images can also be reconstructed. If pre-scan raw data are unavailable, the coil sensitivity estimation concludes here. Otherwise, inspired by ^7^, we further refine the coil sensitivity maps, *C*, using a heuristic alternating gradient descent approach. This refinement alternates between optimizing the reconstructed image **x** and the coil sensitivities *C*, progressively improving the estimated sensitivity maps.

The alternating gradient descent method heuristically minimizes the following objective function:

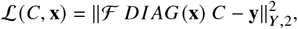

where ℱ is the spatial Fourier transform, and *DIAG* (**x**) is a diagonal matrix containing the values of **x** replicated *nC h* times as follows:

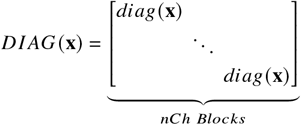

with each block *diag*(**x**) being the diagonal matrix listing the values of **x**. Function ℒ ensures that the forward model—comprising the reconstructed image, coil sensitivity maps, and Fourier transform—matches the acquired *k*-space data **y**. The optimization alternates between the following steps:

1. **Keep** *C* **constant and optimize x:**

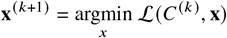 This step refines the reconstructed image using the current coil sensitivity maps.
2. **Keep x constant and optimize** *C*:

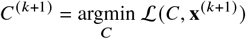 This step updates the coil sensitivity maps based on the current reconstructed image.

To prevent divergence of the two optimized variables (e.g., *x* → *x*^#^/*α* and *C* → *αC*^#^ for some positive scalar *α* → +1), the body coil sensitivity is concatenated with the surface coil sensitivities as an additional channel and is held constant during the optimization process, ensuring that only the coil sensitivities of the surface coils are updated, while the body coil serves as a fixed reference. The concatenated body coil acts as a reference, stabilizing the scale and preventing divergence. To ensure an equilibrated weighting of each surface coil sensitivity and the body coil sensitivity in the iterative refinement, a normalization step, acting on all the images, is performed at the start of the estimation process.

In Monalisa, several functions are provided to estimate coil sensitivity maps, and our documentation offers detailed guidance on how to use these tools^††^. Among these, the mlComputeCoilSensitivity function offers a particularly simple interface for estimating coil sensitivity maps:

`C = mlComputeCoilSensitivity(BCreader, HCreader, optional_arguments)`

The sensitivity estimation produce a MATLAB array of single-precision values with dimensions [*C*_ *fr Size, nCh*]. Here, *nC h* represents the number of receiver-coils, each coil yielding its own sensitivity map, and *C*_ *fr Size* is the frame size of the sensitivity maps during the estimation of *C*, which we usually choose to be much smaller than the image frame size. By estimating *C* at a lower spatial resolution, we reduce the adaptation of the estimated sensitivities to noise in the pre-scan data. This strategy is effective because larger voxels (resulting from lower spatial resolution) contain more signal, thereby increasing the SNR. A common choice is *C*_ *fr Size* = 64 for robust estimation. This low resolution approach also shortens the computation time for the coil sensitivity estimation. At reconstruction time, the coil sensitivity will be interpolated to the size [ *fr Size, nCh*], in order to match the image frame size.

For completeness, we provide the pseudocode of our coil sensitivity estimation in Algorithm 1.

#### 0.0.5 Using multiple body coils helps enhance coil sensitivity homogeneity

In subsubsection 0.0.4, we introduced SIGMA, highlighting its ability to exploit two pairs of body coils rather than relying on a single one. The purpose of this section is to illustrate, with a representative example, the benefit of using multiple body coils to improve the estimation of coil sensitivity.

In Figure 2 , we present a representative example illustrating the inherent inhomogeneity of single body coil images. The last two columns distinctly show that both body coil images exhibit substantial deviations from the RMS image, which serves as a reference for a more uniform image. If the assumption of ^6^ was valid—namely, that the body coils are homogeneous—no variation would be visible. These discrepancies indicate non-uniform sensitivity profiles of the body coils, strongly influenced by their orientation and position within the scanner. This observation underscores the limitations of relying on individual body coils for accurate sensitivity estimation. Constructing a virtual reference coil as a combination of measurements from both body coils offers a more robust solution. In facts, assuming that the inhomogeneities in each body-coil sensitivity are independent and random, averaging their contributions yields to an estimate with reduced variance. This is consistent with the central limit theorem, resulting in a more robust sensitivity map than that obtained from a single body coil.

##### Algorithm 1 *SIGMA* Coil Sensitivity Estimation

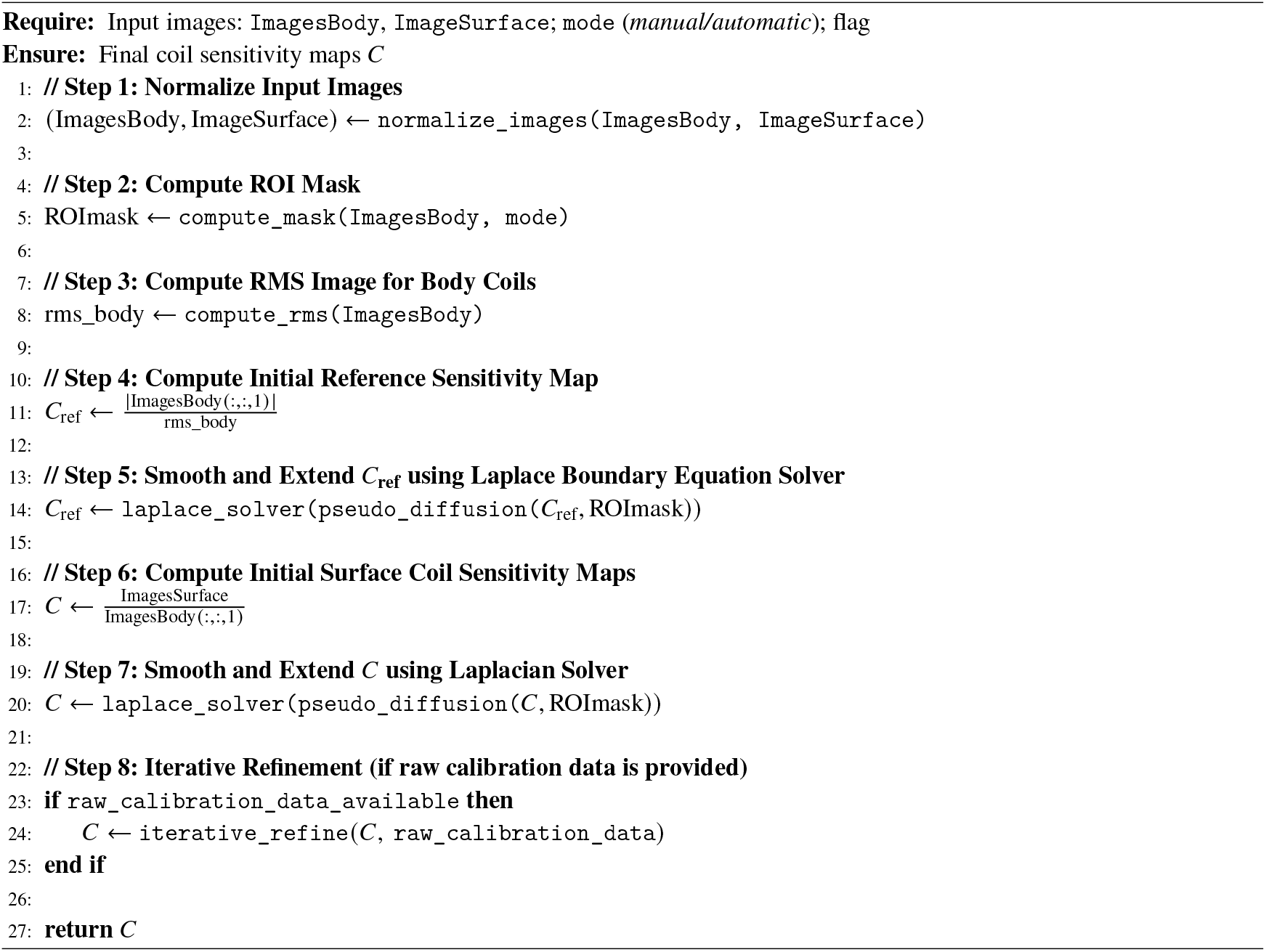

#### 0.0.6 Binning: Flexible Readout Partitioning

In MRI data processing, binning refers to the grouping of readouts into subsets (bins), with each subset specifying the data used for reconstructing a corresponding image frame. In addition to partitioning, binning often involves excluding certain readouts that do not meet specific criteria, such as non-steady-state measurements at the very beginning of the acquisition. For example, while acquiring raw MRI data of the eye with an external eye tracker device, we can use an eye tracker to exclude the k-space points that are affected by motion, or to group in different bins readouts acquired while the eyes was staring at different positions ^16^. Monalisa requires the user to define each of the binning masks *B*_1_,… , *B*_*nB*_ as a logical array, where **nB** is the number of binning masks. Here, **B**_**b**_ denotes the binning mask number *b* and is of size [**nBin**_**b**_, **nLines**] where:

- **nBin**_**b**_ is the number of partitions expressed in binning mask number *b*,
- **nLines** is the total number of lines in the acquisition trajectory, which do not need to be straight lines (e.g. spiral lines). Note that we mask entire k-space lines rather than individual points, as all points on a given line are acquired in a very small time interval.
- **B**_**b**_(**i, j**) is a Boolean value (true or false), indicating whether the *j* -th line contributes to the *i*-th bin of binning mask number *b*.

Each binning mask usually encodes for a different non-spatial dimension. For this reason, if multiple binning masks are used, they are combined using a Cartesian product. As an illustrative example, consider the reconstruction of dynamic CINE images ^17^ using both cardiac and respiratory gating. Suppose we want to resolve six cardiac phases and eight respiratory phases. This setup effectively maps each readout in real time into two distinct non-spatial dimensions: one corresponding to its position within the cardiac cycle, and the other to its position within the respiratory cycle. In this case, two separate binning masks are required—one for the cardiac pseudo-time and one for the respiratory pseudo-time—so we have *nB* = 2. Let us denote the cardiac binning mask as *B*_1_ and the respiratory binning mask as *B*_2_. Then, *nBin*_1_ = 6 and *nBin*_2_ = 8, meaning that *B*_1_ partitions the data into six cardiac bins, and *B*_2_ partitions it into eight respiratory bins. By combining the two masks, the total number of reconstructed frames will be *nBins* = *nBin*_1_ · *nBin*_2_ = 48, as each unique combination of cardiac and respiratory phase defines a single image frame.

**FIGURE 2.**
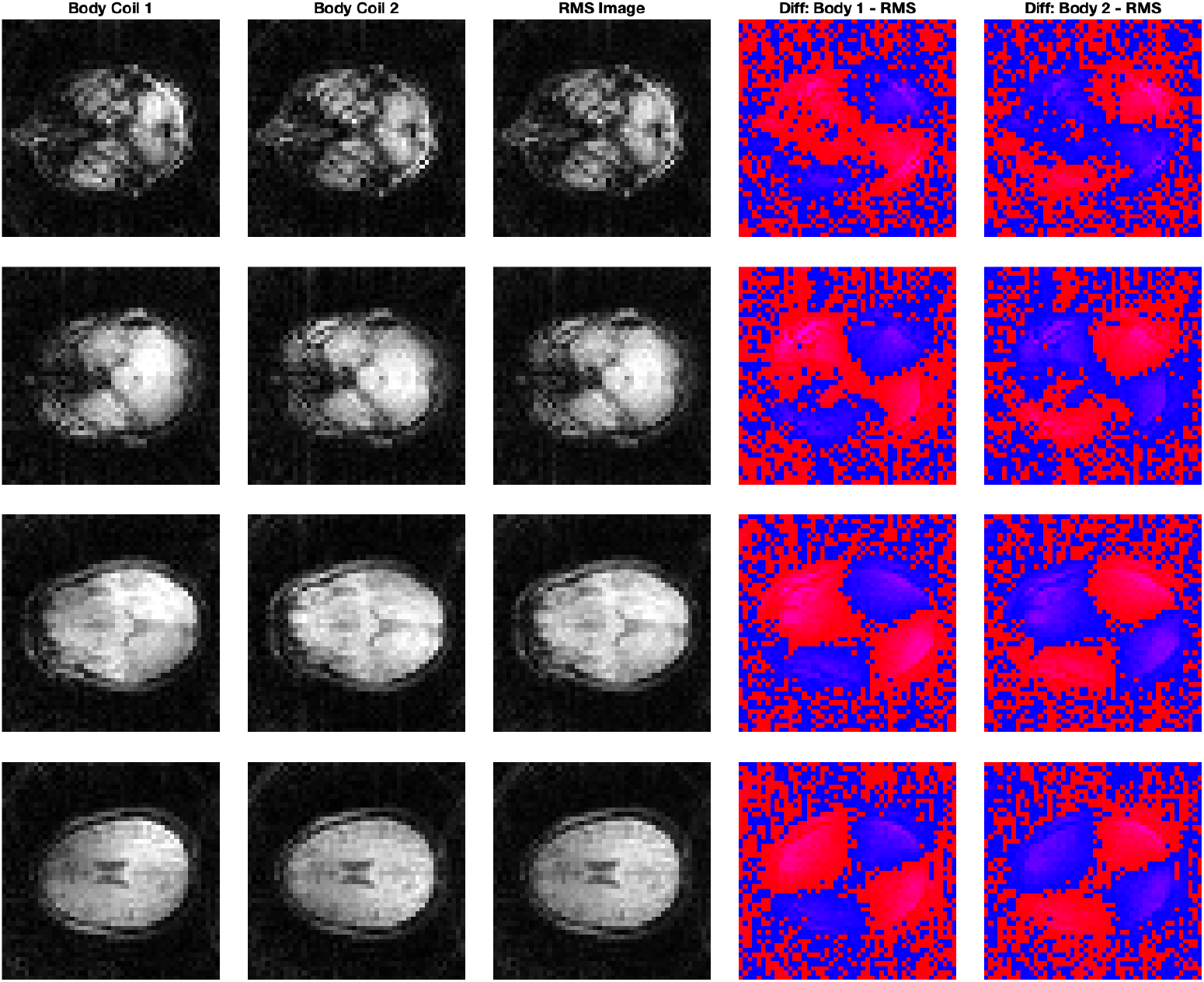
Comparison of individual body coil images and the corresponding RMS image for a single brain calibration scan. Results presented in this image are consistent across all calibration scans. The last two columns reveal significant inhomogeneities in the body coil images relative to the RMS, highlighting the orientation of the body coil in the MRI scanner and their resulting sensitivity variations.

It follows that the total number of resulting bins *nBins* = *nBin*_1_ · … · *nBin*_*nB*_ = *nFr*, because for each bin a corresponding frame will be reconstructed. Since the binning generation is experiment-specific, our documentation provides a few examples to guide the user in implementing their own binning masks^‡‡^.

#### 0.0.7 The Seed and the Mitosius: A Standardized Data Structure to Store the Result of the Reconstruction Preparation

In Monalisa, reconstructions are performed in two main sequential steps: a preparation step, followed by the reconstruction itself. Users are free to design the preparation step however they wish, as long as its output includes all the arrays and parameters required to run the reconstruction function.

Nevertheless, this section gives an overview of how to use the standardized data structure defined for Monalisa, that we highly recommend. This structure consists of two folders: the “seed” and the “mitosius”, as illustrated in Figure 1 . The seed folder, whose name can be freely chosen by the user, serves as the main reconstruction data container. It contains all the necessary inputs to start the reconstruction process and all the resulting outputs are written within it. Typically, the seed is the folder transferred to an High-Performance Computing (HPC) system to perform high-dimensional reconstructions. The mitosius is a subfolder of the seed and its name cannot be changed, it is part of the standard. The mitosius folder organizes the data *y*, trajectory *t* and volume elements *ve* for each bin in a separate subfolder, referred to as a “cell”. For single-frame reconstructions, the mitosius folder contains only one cell. For multi-frame reconstructions, the mitosius folder contains *nFr* subfolders, each containing *y, t*, and *ve* of a single bin.

The estimated coil sensitivity maps are not stored within mitosius. Instead, they must be placed in a subfolder named “C” within the seed. In addition, the binning masks are not required to include all the acquired data. When this occurs, the mitosius reduces the amount of data copied to the reconstruction algorithm. This is an advantage when working with HPC, since the data transfer can be time-consuming.

Creating the seed and mitosius completes the preparation step. To create the seed, create a new folder with the name of your choice. Next, create the mitosius within the seed, to do that ensure first that you have access to:

- **Acquired raw data y**: It can be simply read from Siemens raw data or ISMRMRD files using Monalisa’s readers as previously described in subsubsection 0.0.1. It contains the totality of the acquired raw data.
- **Acquisition Trajectory t**: Computed as previously described. If you did not compute it yet, you are now at the right step to compute it. It is the raw trajectory that was used during the acquisition. It therefore contains the totality of the trajectory points.
- **Coil sensitivity map C**: Computed as previously described in subsubsection 0.0.4.
- **Binning Masks B**_**1**_,… , **B**_**nB**_: Computed as previously described in subsubsection 0.0.6. *nB* is the number of binning masks.

For single-frame reconstructions, volume elements *ve* for *t* are computed as described in subsubsection 0.0.3. In case of multi-frame reconstruction, the user can run the “mitosis” function before evaluating the volume elements. This function builds the different data bins and trajectory bins according to the binning mask(s) given as input. The mitosis function is invoked using the following syntax:

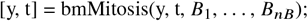

This function updates *y* and *t*, converting them into MATLAB cell arrays in which each cell corresponds to a specific bin. Conceptually, it partitions the original data arrays into smaller units, that are named cells (this is the origin of the name mitosis). The mitosis function transforms the complete **acquired data** into the **reconstruction data**, and the complete **acquisition trajectory** into the **reconstruction trajectory**.

Once the mitosis is performed, volume elements for each cell are computed as described in subsubsection 0.0.3 using *t* and stored in a corresponding cell-array *ve*. Note that each binning mask must be of size [*nBin*_*b*_, *nLines*] where *nBin*_*b*_ is the number of bins in the binning mask number *b* and *nLines* is the number of total acquired lines. After the mitosis function, *y* and *t* are cell arrays of size [*nBin*_1_, …, *nBin*_*nB*_], where each cell contains an array with dimensions determined by the contents of the corresponding bin. The mitosis function supports any number of binning mask. However, Monalisa’s reconstructions currently support only one-dimensional cell-arrays (that we call “chains”, *nB* = 1) and two-dimensional cell-arrays (that we call “sheets”, *nB* = 2).

At this point, it is recommended to perform a rescaling of the raw data *y* by a positive value that we will call *v*_*scale*_. To choose *v*_*scale*_, we suggest that the user perform a preliminary gridded (zero-padded) reconstruction with the function “bmMathilda” if the data are non-Cartesian (or “Nasha_partialCartesian” for Cartesian data), display the magnitude image, draw a ROI, compute the average magnitude of the image in that ROI, and set *v*_*scale*_ to be equal to such value. This operation ensures that the reconstructed image will have an average value close to 1 in the ROI selected during the rescaling process. We observe that if *y* contains multiple cells, rescaling implies dividing the content of each cell by the same value *v*_*scale*_. Additional information and instructions are available in our documentation^§§^.

Finally, permute each array *y*{*i*} so that its size is [*nPt*{*i*}, *nCh*], where *nPt*{*i*} is the number of trajectory points selected by bin number *i*. By running the function bmCreateMitosius^¶¶^ , the mitosius folder with its content will be saved on the disk: in each subfolder of the mitosius, you will find the arrays *y, t* and *ve*.

This terminates the preparation step. You are now ready to reconstruct your images. If you are working with an HPC cluster, the only folder you need to transfer is the seed, which contains the mitosius, your coil sensitivity estimates and other reconstruction parameters that you can load just before the call of the reconstruction function in the script running on the HPC.

### Determination of the Number of Iterations

In this section we show how we selected the number of iterations, that was set to 160. In Monalisa, which allows saving intermediate results, we monitored the evolution of reconstruction metrics throughout the iterative process. This enabled us to assess convergence directly. Figure 3 illustrates a representative example for the *l*1-regularized reconstruction of the eye image, demonstrating that convergence indeed occurs before 160 iterations.

**FIGURE 3.**
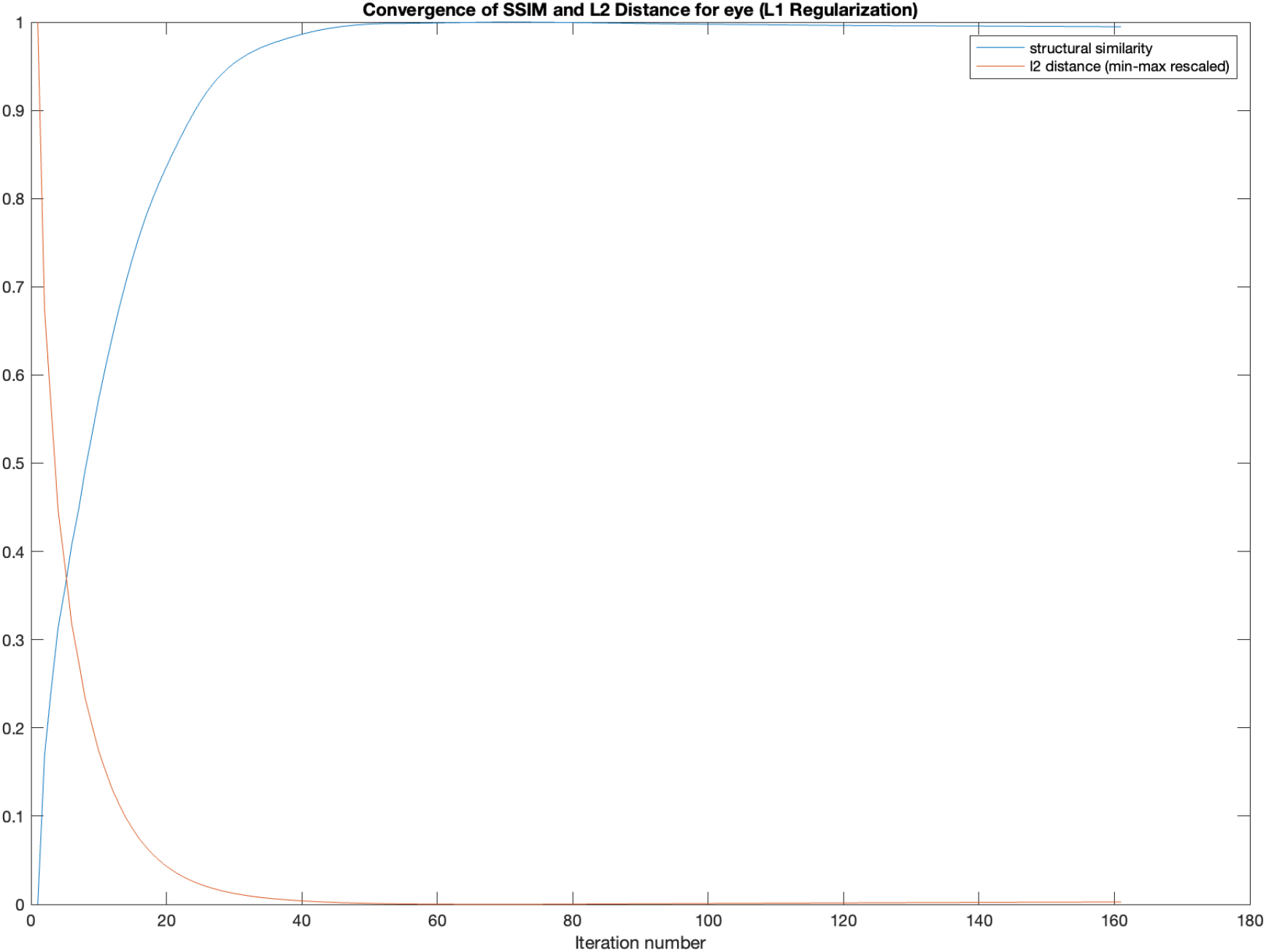
This figure shows the convergence of Structural Similarity Index (SSIM) and *l*_2_ distance over iterations for an iterative reconstruction. In particular, this example corresponds to a Monalisa reconstruction using *l*_1_ regularization on the eye synthetic data. The curves demonstrate that both metrics stabilize, indicating convergence of the reconstruction process.

**FIGURE 4.**
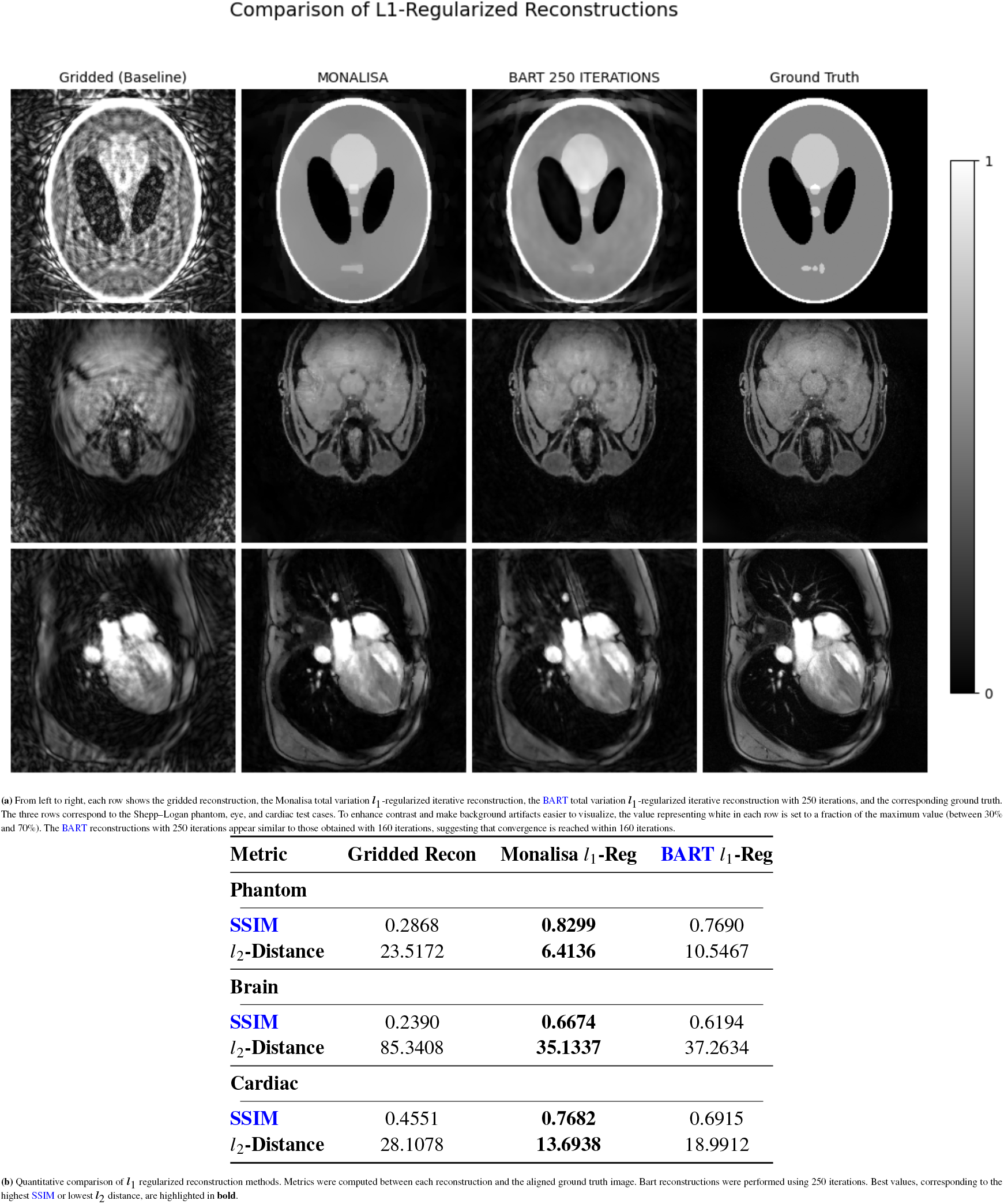
Comparison of *l*_1_-regularized reconstruction results using visual and quantitative metrics. Bart reconstructions were performed using 250 iterations.

**FIGURE 5.**
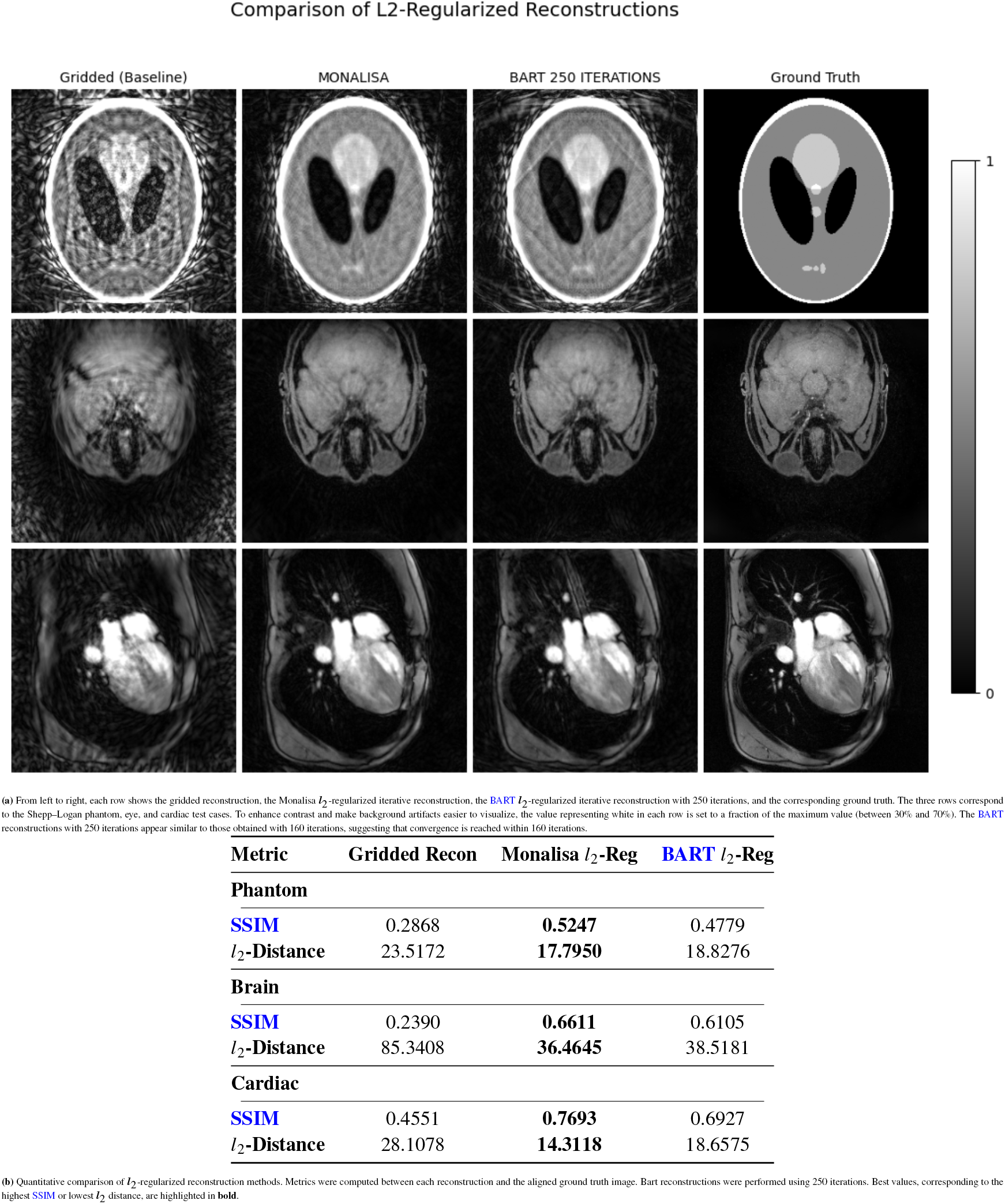
Comparison of *l*_2_-regularized reconstruction results using visual and quantitative metrics. Bart reconstructions were performed using 250 iterations.

**FIGURE 6.**
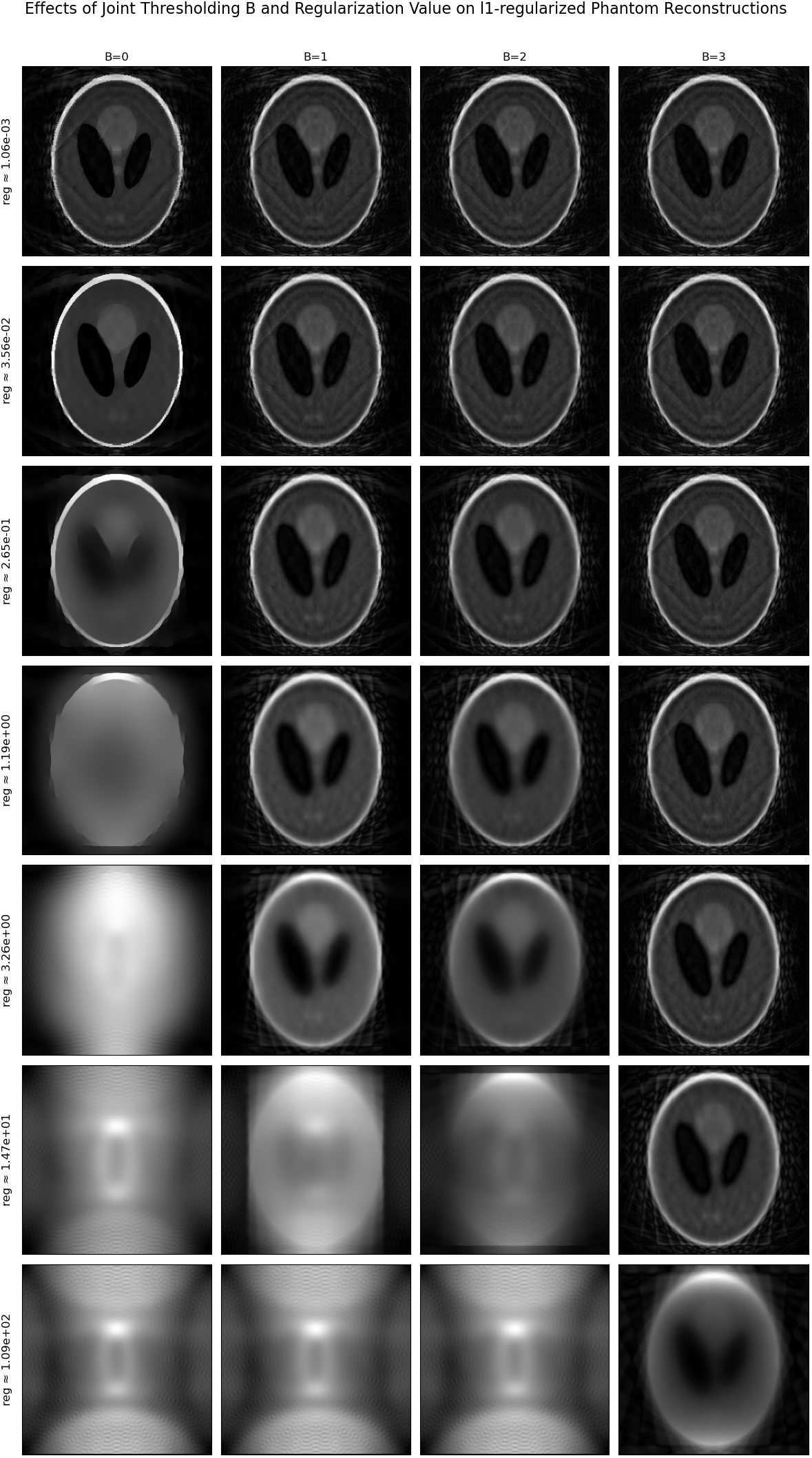
Effects of joint thresholding parameter *B* and regularization value on total variation *l*_1_-regularized reconstruction quality for BART framework using generalized regularization syntax. Our experiments indicate that changing *B* shifts the optimal regularization value and best image quality is observed using B=0.

For Berkeley Advanced Reconstruction Toolbox (BART), intermediate results could not be saved. Since running 160 reconstruction was unreasonable, to verify convergence, we tested multiple iteration counts. As a proof of convergence, we include results identical to those previously presented, with the only difference being that the BART reconstruction was performed with 250 iterations instead of 160. For the *l*_1_-regularized case, the differences are minimal but visible in the table. For the *l*_2_-regularized case, the changes are negligible, with differences less than 10^–4^.

### Joint thresholding flag for BART’s *ℓ*_1_-regularization

In BART, total variation *4*_1_-regularization in the image domain is invoked using the generalized regularization syntax:

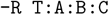

where *A* specifies transform flags, *B* is the *joint thresholding* flag, and *C* is the regularization value. In the documentation, no description of joint thresholding and its effect is provided.

We assume that joint thresholding modifies the definition of the regularization term *R*(*x*) in.

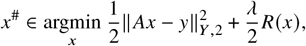

possibly by linking thresholding decisions across dimensions, channels, or other image components.

Through empirical testing and reverse engineering, we found that only the two least significant bits of *B* influence the reconstruction, yielding exactly four distinct output modes. This is plausible, and possibly similar to the bitmap masking adopted by BART. This suggests the presence of two independent thresholding mechanisms or dimensions, with *B* determining whether none, one, or both are applied.

## Footnotes

∗ See implementation here

† See implementation here

‡ See implementation here

§ mapVBVD. As stated in our license, these functions are not part of the toolbox and are located in the subfolder /monalisa/third_part/twix_for_monalisa/.

¶ Be aware that in Siemens scanners, if you input a given FOV in the readout direction, the true FOV will be twice the input value entered.

\# See: How to use custom trajectory

∥ See ^3^ for gridded reconstructions, see here for iterative reconstructions.

∗∗ See: Volume Element Computation

†† See: How to compute the coil sensitivity

‡‡ See: How to bin data

§§ See: Normalizing data

¶¶ Source code of bmCreateMitosius

